# Roles of motion perception and visual acuity for driving hazard perception

**DOI:** 10.1101/355958

**Authors:** Mojtaba Moharrer, Xiaolan Tang, Gang Luo

## Abstract

**PURPOSE:** There are many visually impaired people who can drive legally with bioptic telescope. Drawing on the experience of drivers with reduced vision, this study investigated the role of motion perception and visual acuity in driving, under simulated low visual acuity.

**METHODS:** Twenty normally sighted participants took part in a driving hazard perception (HP) test, in four different conditions: with/without motion interruption and with/without simulated low visual acuity. In interrupted motion conditions a mask frame was inserted between every frame of the driving videos. In simulated low visual acuity conditions, participants wore glasses with diffusing filters that lowered their visual acuity to 20/120 on average. Participants’ response time, hazard detection rates, and HP scores, which combined response time and detection rate, were compared.

**RESULTS:** Repeated measure ANOVA revealed that the HP scores significantly declined from 20.46 to 16.82 due to the motion mask (F_(1,19)_ = 9.343, p = 0.006). However, simulated low visual acuity did not affect HP scores (F(_1,19_) = 1.807, p = 0.195). The interaction between vision and mask was not significant (F(_1,19_) = 1. 295, p = 0.269). The decline in score was mostly due to significant decrease in detection rate, from 0.80 to 0.64, due to the motion mask (F(_1,19_) = 16.686, p = 0.001).

**CONCLUSIONS:** In this experimental setting, human observers relied largely on motion information for detecting driving hazards, rather than high visual acuity. This finding might help explain how visually impaired drivers can compensate for their impaired vision during driving.

## Introduction

Visual cues provide a major source of information when driving. Historically visual acuity has been used as the main visual test for assessing drivers’ visual fitness, however, it is not clear which aspects of vision are critical for safe driving as studies have shown different, and at times conflicting, results ^1^. Owsley and McGwin Jr ^2^ comprehensive review of the existing research on vision and driving showed the different aspects of vision found to be influential in safe driving. They concluded that “visual acuity is related to certain aspects of driving performance”, however, “*at best, very weakly linked to driver safety*” ^2^.

In addition to visual acuity, studies have tried to explore the relationship between safe driving and visual functions, including contrast sensitivity ^3–6^, visual field ^7–9^, and motion perception ^10,11^. These studies also showed that there is more to the relationship between vision and driving than visual acuity alone.

A high number of the studies into vision and driving focus on normally sighted drivers, a comprehensive literature review of these studies is provided by Owsley and McGwin Jr ^2^ and Owsley, Wood and McGwin ^1^. The use of normally sighted drivers offers opportunities to look at only a limited range of visual functions for investigation of role of vision on driving. In order to gain a more comprehensive understanding of the relationship between vision and driving and the impact vision has on driving, research is needed on a wide range of visual functions. To achieve this, there is a need to include drivers with visual impairment in the studies investigating vision and driving.

Many states in the United States, as well as the Netherlands and Quebec, permit drivers with low levels of visual acuity, as low as 20/200 in some cases, to drive with the assistance of a visual aid called bioptic telescope ^12,13^. Using varying methods, studies have investigated the safety performance of bioptic users; some found bioptic telescopes users to be safe drivers ^14–17^, whilst others found bioptic users to be slightly less safe than normally sighted drivers ^18–20^.

A retrospective study of the relationship between vision and bipotic drivers’ perfromance in road test^21^, showed that the hours of training, not any visual factors, were the predictive of road testing outcome. Also, Dougherty, Flom, Bullimore and Raasch ^15^ study on traffic collisions of a group of bioptic drivers found that more driving experience prior to the use of bioptic telescope correlated with lower possiblity of being involved in road crashes. Their study ^15^ did not find any significant relationships between vision (visual acuity and contrast sensitivity) and road collisions. Nonetheless, bioptic users who have driven safely for decades provide possibilities for targeted research on the relationship between vision and driving. These drivers’ low level of visual acuity sets them apart from the drivers with conventional levels of visual acuity. Bioptic drivers use the telescope mainly for navigation and reading the signs ^22,23^, which means that they use their bioptic telescope for only a small proportion of total driving time, adhering to the bioptic driving guidelines which addresses concern that use of the bioptic telescope may block the field of view ^24^. Luo and Peli ^12^ recorded the actual driving of two bioptic drivers and found that their bioptic use was as low as 1%, meaning that for 99% of the time these drivers drove with their low visual acuity. The purpose of this current study is to investigate whether low visual acuity hampers drivers’ ability to perceive driving hazards.

## Visual Acuity, Motion Perception and Driving

Visual acuity is the visual system’s capacity for resolving spatial information, “the ability of the eye to see fine detail” ^25^. While seeing fine details is important for many daily activities, motion provides an additional source of information for perception of objects ^26^. Studies have shown that low spatial frequency information is sufficient for perception of moving objects ^27, 28^ and consequently, motion perception can help scene comprehension even when the images are blurred.

In their study, Saunders, Bex, Rose and Woods ^29^, designed a method for measuring information acquisition and applied it to the perception of movie clips. Participants used varying spherical defocus lenses to simulate reduced levels of visual acuity, and they were then asked to give a verbal description of some short movie clips. Their responses were compared to a database of natural language responses made by normally sighted participants following viewings of the same video clips, transcribed using Amazon Mechanical Turk. The study found that information acquisition with the visual acuity conditions of as low as 20/125 did not differ significantly from the 20/20 condition.

Pan and Bingham ^30^, studied motion perception using blurry images extracted from daily activities, such as playing basketball, bowling and washing the dishes. Twenty images were extracted from the videos and were presented in five conditions, including, (1) three random static single frames, (2) all twenty frames with the motion masks playing in the loop, (3) all twenty frames in order without the motion masks playing in the loop ^30^. In the second condition, where the motion mask was presented, the duration of the image was 0.5 seconds and the duration of the mask was 2 seconds. The first two conditions, static image and with the motion mask, showed significantly lower correct events perceived compared to the without the motion condition. Their results showed that “motion could calibrate blurred images to allow subsequent accurate perception of events despite limited image information” ^30^.

Studies have examined the role of motion perception in different aspects of driving ^10,11,31–34^. Hazard Perception (HP), the ability to anticipate traffic, is considered one of the main safety related driving skill ^35^. The HP test, explained in detailed in Method section, is a part of the driving license test in the UK and Australia ^36^. Researchers have widely used HP in research on driving ^10,11,37^. Wilkins, Gray, Gaska and Winterbottom ^11^ investigation of the relationship between motion perception and driving with a group of normally sighted participants used motion perception tests that included 2D motion-defined letter identification, 3D motion in depth sensitivity, and dynamic visual acuity tests. The driving tests were conducted using a driving simulator and included HP and emergency brake tests.

Wilkins, Gray, Gaska and Winterbottom ^11^ found that higher 3D motion perception was correlated with better emergency brake performance, and higher dynamic visual acuity was significantly correlated with better HP test performance. The results did not show a significant relationship between HP and motion perception tests, which is in contradiction with the findings of other studies ^10,32,38^. Wilkins, Gray, Gaska and Winterbottom ^11^ suggested two possible explanations for their finding: first, the complex design of the motion test employed required participants to indicate the letter they were presented with, rather than solely identifying the direction of the motion. Second, the limited driving experience of participants in their study, which can lead to reduced HP performance. In addition, Wilkins, Gray, Gaska and Winterbottom ^11^ performed a set of training, with the aim of improving participants’ motion perception and exploring its impact on emergency braking performance. For the training section, participants were asked to take the same three tests of the first section, 2D motion-defined letter identification, 3D motion in depth sensitivity, and dynamic visual acuity test, over a six week period and where auditory feedback were presented. Results of the study showed a significant improvement in brake reaction time for the participants that took part in the training while the control group did not show a significant change.

In a research by Lacherez, Au and Wood ^10^ the association between motion perception and HP was studied. A set of visual measures were examined: Static visual acuity, Pelli-Robson letter contrast sensitivity, Visual fields, and two motion perception tests Random dot kinematogram ^38,39^ and drifting Gabor patch ^40^. Lacherez, Au and Wood ^10^ found significant relationship between HP and both motion perception measures they used, where a better HP performance correlates with better motion perception measures, Random dot kinematogram and drifting Gabor patch. They also found that better HP performance was correlated with better visual acuity. Lacherez, Au and Wood ^10^ conclude that “motion perception plays an important role in the visual perception of driving-relevant hazards” and suggest a need for further exploration of the causes of reduced motion perception.

Wood, Black, Mallon, Kwan and Owsley ^34^ investigation of the visual characteristics of age-related macular degeneration (AMD) driving population looked at 33 AMD participants and 50 age-matched control participants, all above 65 years old. The different visual characteristics that were tested include visual acuity, contrast sensitivity, visual fields, and motion sensitivity, along with participants’ on-road driving performance. Participants’ on-road driving performance was rated by an occupational therapist who was unaware of drivers’ visual status. Results of Wood, Black, Mallon, Kwan and Owsley ^34^ study showed that motion sensitivity was the only visual measure significantly associated with driving safety of AMD drivers.

Building on past research, this current study argues that motion perception is essential for safe driving. Unlike the previous studies that analyzed the relationship between HP and motion perception ^10, 11^, the aim of this current study is to find the causality of motion perception and HP by removing/reducing the motion perception, similar to the approach used by Pan and Bingham ^30^. In addition, the current study explores the causality of visual acuity and motion perception within the same individuals, overcoming individual differences when comparing between groups.

## Method

### Hazard Perception

The HP test is widely used in research into safe driving and is a part of the driving license issuance procedure in the UK, in which a total of 15 hazards are introduced and participants are directed to press a key as soon as they see a “potential hazard” or a “building hazard” ^41^.

There are two direct outcomes from HP test: first, whether the participant notices the hazard and responds to it, and second, the participants’ reaction time, (how quickly the participant responded to the observed hazard). Using the first outcome detection rate is calculated, showing the percentage of hazards detected while the hazard was still present and before the driver in the video reacted. The second outcome of the HP test is the reaction time, which shows the mean reaction time of all hazards detected from the first moment of appearance on the screen.

In order to have a comprehensive measure that combines both reaction time and detection rate, the scoring system currently used for the UK’s driving license examination was utilized for measuring HP test. In this scoring system each of the fifteen hazards is allotted five points, making a total score of 75 points. Scores are calculated based on the response time to each presented hazard. The total time from the moment that hazard appears until the time the driver in the recorded video reacts or the hazard eliminates is measured and scaled from zero (too late or no detection) to five (very fast detection). The score then is calculated based on the response time in that scale. The benefit of using HP score is that it includes both reaction time and detection rate elements while measuring overall performance.

### Stimuli

The design of the study method suggests that an important part of motion perception is the continuity of the presented images, optic flow. In order to identify the role of motion perception in HP, a method, similar to Pan and Bingham ^30^ was used in this study in order to remove/reduce the perception of the motion in HP tests. This was accomplished through the interruption of the optic flow in presentation of continuous motion. The method involved a mask image inserted between the frames of HP videos. In their study, Pan and Bingham ^30^, used a white mask for 2 seconds, and event images were shown for 0.5 seconds. It can be argued that the duration of mask was so long that observers essentially viewed stationary snapshots. To address this type of critique, this study used a much shorter mask duration. In addition, during the pilot phase, three different blank color frames (black, gray, and white) and an average image, which were created based on the average Fourier spectrum of 500 frames randomly extracted from the HP test videos, were tested. The mask created has the same characteristics of spatial frequency as the original images, but does not include any meaningful scene information, except for the overall brightness pattern of driving scenes. The feedback received from participants revealed that the blank color frames were very distracting because of sudden brightness change. In response to this feedback, the average image was used as the mask.

The original frame rate of HP videos was 25 frames per second. In order to accommodate the insertion of mask frame and also maintain the video duration same as the originals, from every ten frames one frame was extracted and presented to participants. In no-mask conditions, each frame was shown for 0.4 seconds. In with-mask conditions, the duration of image and mask remained the same 0.4 seconds. To test the impact of mask duration on HP performance, two mask durations, 0.1 seconds and 0.3 seconds, were used. For thirteen participants, in masked condition, the image was shown for 0.1 seconds while the mask was shown for 0.3 seconds. For the remaining seven participants, in the masked condition, the image was shown for 0.3 seconds while the mask was shown for 0.1 seconds. For a better explanation, these three timings are shown in Figure 1.

**Figure 1.**
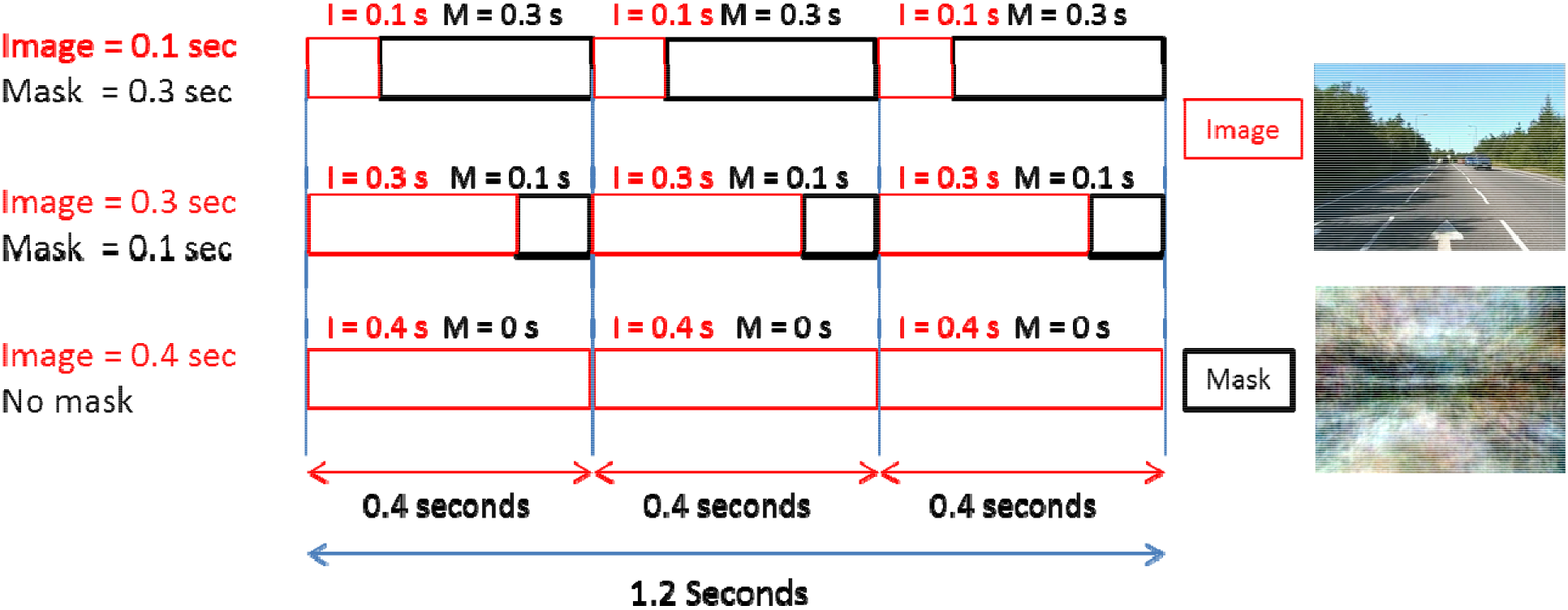
- Three designs of Image and Mask timings. From every ten frames one frame was extracted and presented to participants. The total duration of all videos remain the same as the original videos. Two mask durations, 0.1 seconds and 0.3 seconds, were tested in this study.

### Testing conditions

Twenty normally sighted participants, ten male and ten female, were recruited for this study. To simulate reduced visual acuity, participants wore a pair of glasses with the plano lenses covered with two 0.1 diffusing foils (Bangerter occlusion foils, Ryser Optik AG, St. Gallen, Switzerland). On average, the diffuser reduced visual acuity to 20/120 in normally sighted people. The design of the experiment was a two-by-two combination of vision (with/without diffuser) and motion conditions (with/without the motion mask). The order of the four conditions was counterbalanced across participants.

To ensure accuracy in the presentation of the HP videos, a program was created using PsychToolbox. Forty HP videos were shown in blocks of five (ten videos for each condition) with each video lasting approximately sixty seconds and containing one hazard. The videos were taken from a commercial training DVD used for driving license test in the UK (Driving Test Success Hazard Perception: Imagitech Ltd, Swansea, UK). The HP videos were played on a 23” Asus monitor, model VX238H-W, with screen resolution of 1920*1080 and frequency of 60 Hz. The HP videos were 786*576, presented with a viewing distance of 60 cm.

Participants were first briefed on the HP test and what they were expected to do and were then asked to watch five sample HP videos, which included different types of potential hazards. In order to confirm participants’ understanding of the task expectations, participants were asked, during the practice, to click the mouse when the hazards appeared and to verbally explain the hazard they observed.

This study was approved by Massachusetts Eye and Ear’s Institutional Review Board and was conducted in accordance with tenets of the Declaration of Helsinki, with informed consent being obtained from the research participants.

### Statistical analysis

For the analysis of the results, there were two independent variables, mask and diffuser, and three dependent variables, Detection rate (between zero to one), Reaction time (in seconds) and HP scores (out of 75). Repeated measure ANOVAs were used to analyze the effects of mask (interruption in motion) and diffuser (simulated low vision) on each of the dependent measures. A *p* value of <0.05 was considered as statistically significant.

### Results

Figure 2 shows the HP detection rate and reaction time results. It can be seen that the mask significantly reduced the detection rates (F(_1,19_) = 16.686, p = 0.001), from 0.803 to 0.640. The use of diffuser also significantly reduced detection rates (F(_1,19_) = 4.643, p = 0.044), from 0.765 to 0.677. The interaction of diffuser and mask was not significant, F(1, 19) = 2.451, p = 0.134.

The repeated measure ANOVA did not find significant effect of motion perception on reaction time, (F(_1,19_) = 1.789, p = 0.197), nor did the diffuser, (F(_1,19_) = 0.761, p = 0.394). The interaction between diffuser and the motion mask was not significant either (F(_1,19_) =0.271, p = 0.609).

**Figure 2.**
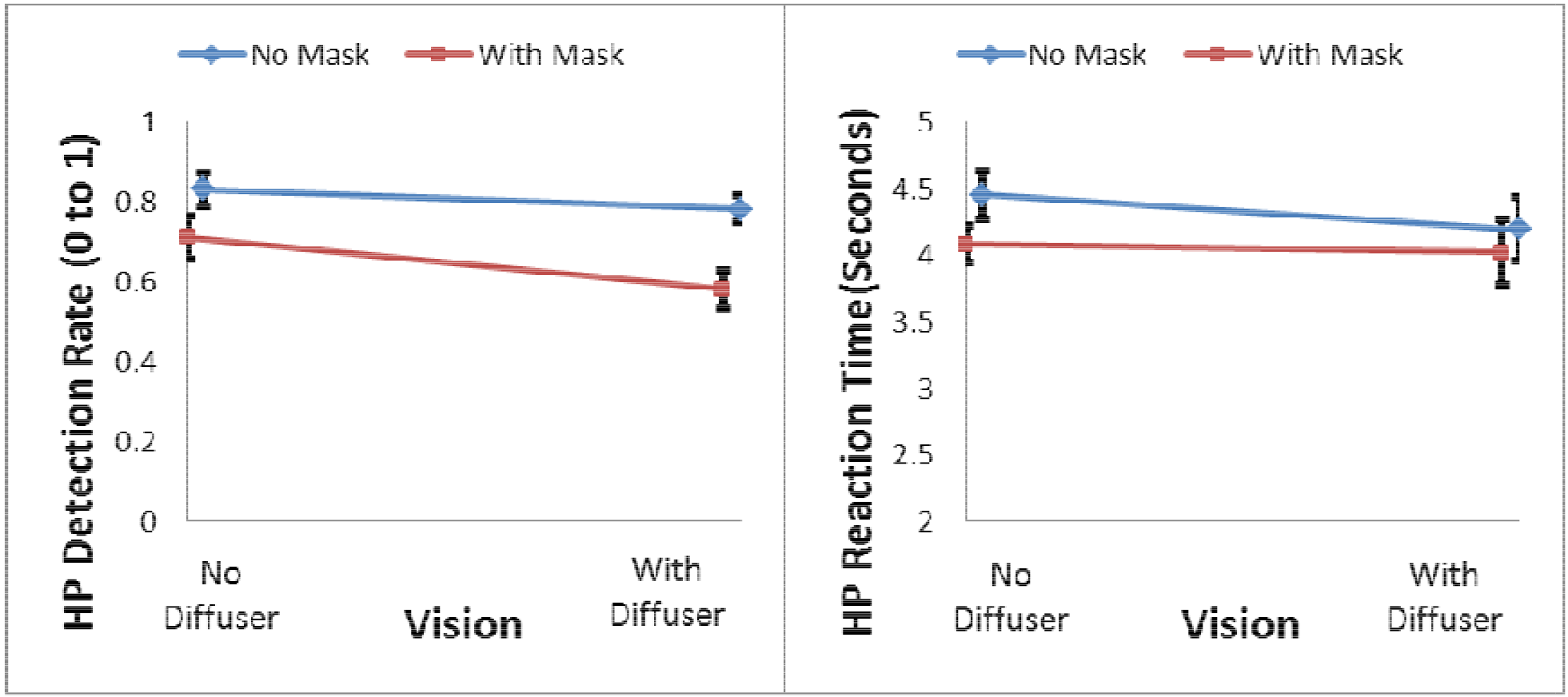
– HP detection rate results (left figure) – Mask shows a strong significant effect (p= 0.001), while diffuser shows a borderline impact (p=0.044). HP reaction time results (right figure) – Mask did not show a significant impact (p = 0.197). Diffuser did not have significant effect either (p = 0.394) – Error Bars: Standard error of the mean

While the diffuser’s effect on reaction time was not significant, when the motion mask was not used, the average reaction time with diffuser (low visual acuity) was shorter than without it (normal vision). Considering the detection rate dropped due to use of diffuser, a further analysis was conducted to determine whether this phenomenon was due to a little too late detection. In other words, when the diffuser was used, could some hazards be detected after the defined hazard response time window, and consequently only easy hazards were considered for reaction time calculation, resulting in shorter average reaction time?

The hazard windows in all videos were extended by 0.4 seconds, to include slightly later recorded responses to the hazards. Following this change, another set of analyses were conducted on the results to include “later” responses. As shown in Figure 3, detection rate was not affected by the diffuser, (F_(1,19)_ = 3.717, p = 0.069), while the impact of mask remains significant, (F_(1,19)_ = 19.086, p < 0.001). As for the reaction time, nether the motion mask (F_(1,19)_ = 1.582, p = 0.224) nor diffuser (F_(1,19)_ = 0.212, p = 0.651) had significant effect.

**Figure 3.**
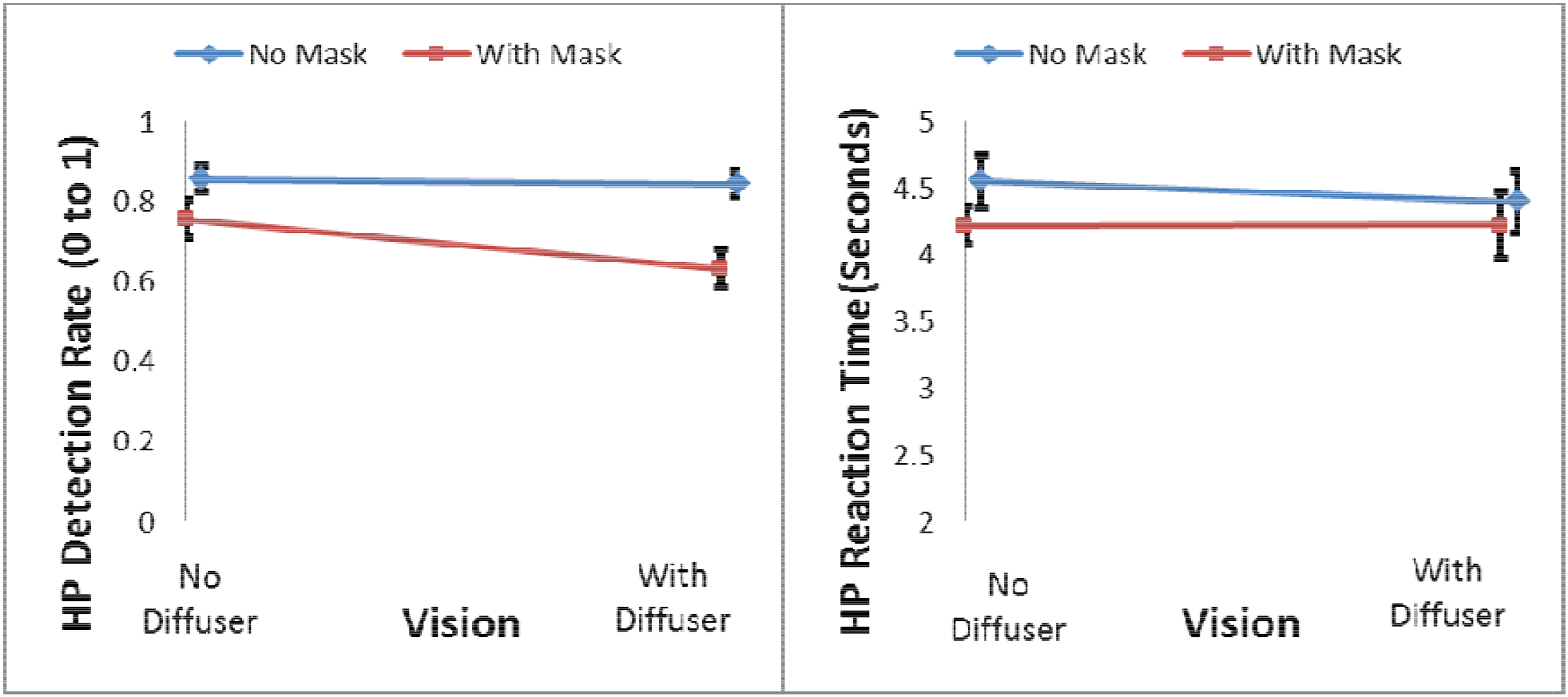
– HP detection rate results with 0.4 seconds delay (left figure) – Mask shows a strong significant effect with p< 0.001 while, diffuser does not show significant change (p=0.069). HP reaction time results with 0.4 seconds delay (right figure) – Mask did not show a significant impact (p = 0.224). Diffuser did not show a significant impact either (p = 0.651) – Error Bars: Standard error of the mean.

Figure 4 shows the results of HP scores, which combine detection rate and reaction time according to HP testing rules used in the UK licensing system^36^. The motion mask significantly reduced HP score from 20.5 to 16.8, (F_(1,19)_ = 9.343, p = 0.006). Meanwhile, the use of diffuser did not have a significant impact on HP performance, (F_(1,19)_ = 1.807, p = 0.195). The interaction between diffuser and the motion mask was not significant (F_(1,19)_ = 1.295, p = 0.269). HP scores were re-analyzed using the 0.4 seconds extended threshold, similar to detection rate and reaction time results, to find the impact of late responses. Similar to initial threshold, only mask showed significant impact on HP score (F_(1,19)_ = 10.731, p = 0.004).

**Figure 4.**
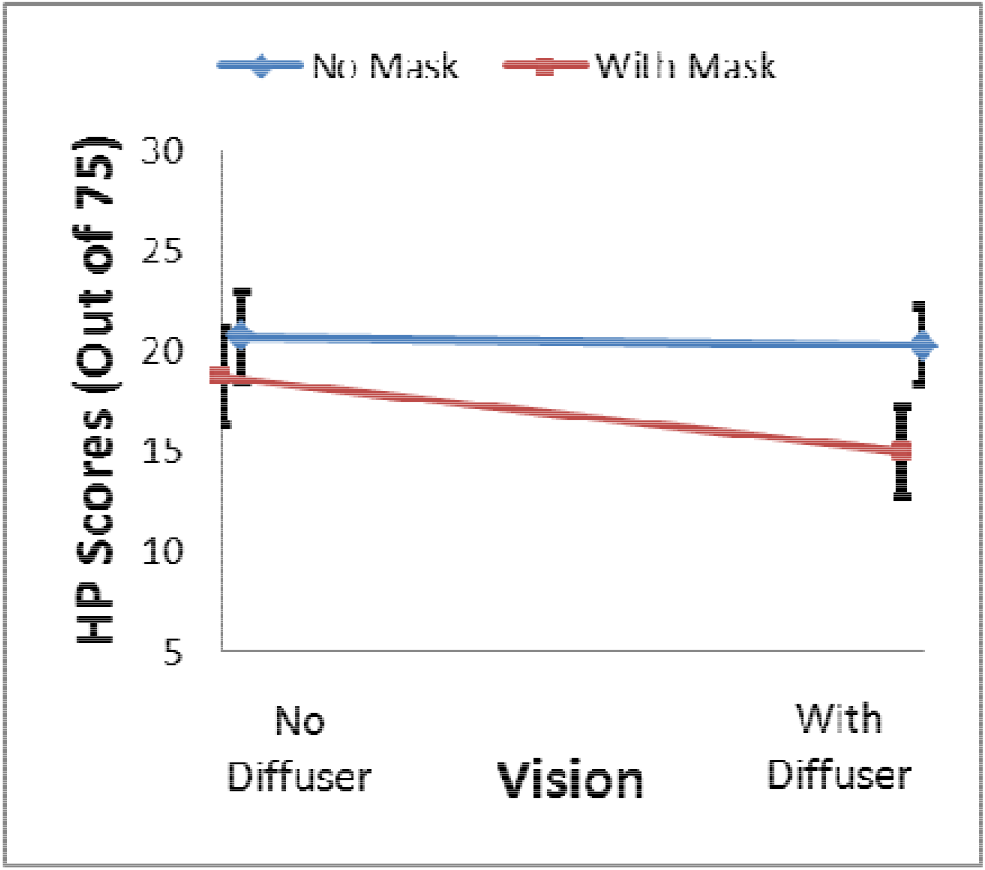
– HP score results – Mask shows a strong significant effect with p= 0.006, while, diffuser does not show a significant impact (p=0.195) – Error Bars: Standard error of the mean

Mask duration (0.1 or 0.3 second) was included as a between-subject factor in repeated measure ANOVA of HP score. The analysis did not find significant effect of mask duration (F_(1,18)_ = 0.677, p=0.421). With this between-subject factor included, the effect of mask still remained significant (F_(1,18)_ = 6.991, p = 0.016) and diffuser was not significant (F=0.682, p=0.420).

The driving images in this study were effectively presented at 2.5Hz, as only one out of 10 frames was shown to participants. To investigate the effect of such a low frame rate of video stimuli, results of this study were also compared to the findings of our previous HP study which used full 25Hz videos ^24^. Results of no-mask-with-diffuser in this study were compared to that of visually impaired participants in previous study, and the results of no-mask-no-diffuser condition of this study were compared to that of normally sighted participants in previous study. HP scores in this study were significantly lower than that in previous study (p <0.002) for both conditions. Results of reaction time also were significantly larger in this study as compared to previous study (p = 0.013). However, the detection rates in this study were not different from previous study (p>0.34).

## Discussion

Building on past research findings on motion perception and driving, this study was designed to explore the causal relationship between motion perception and driving through a HP test. Past research ^10,11,33^have, directly or indirectly, investigated the relationship and correlation between HP and motion perception. Performances of HP and motion perception were measured in different paradigms. For instance, Wilkins, Gray, Gaska and Winterbottom ^11^ examined the correlation between participants’ HP performance and 2D motion-defined letter and 3D motion in depth sensitivity tests, and ^10^ examined the association between HP and two motion perception tests, Random dot kinematogram and drifting Gabor patch. It can be argued that the findings of correlation between the two do not necessarily prove existence of causation. In this study, we directly manipulated motion in the same HP test rather than correlating separated HP test and motion test. We also manipulated visual acuity together with motion to further test their roles in driving hazard perception. In so doing, each participant took the HP test in all four conditions, with and without mask and with and without diffuser.

In order to look at the HP performance of participants, their HP score were used as the main outcome, which combines the two direct outcomes of HP test, detection rate and reaction time. Results of this study showed that the interruption of motion, through mask, resulted in highly significant drop in HP score of participants. At the same time, the reduction in visual acuity, through diffuser, did not show a significant change in HP score of research participants. The impact of interruption of motion perception through mask is in line with previous findings by Pan and Bingham ^30^ that showed the motion mask significantly reduces the perception of events.

Understanding how the low spatial frequency information is used by the human visual system in motion perception might help to explain why visual acuity did not affect HP in this study. Ramachandran, Ginsburg and Anstis ^27^ argued that perceived movement is determined primarily by low spatial frequencies. Tadin, Nyquist, Lusk, Corn and Lappin ^42^ showed that there were no adverse effects of visual acuity loss on motion perception for low spatial frequency stimuli. Our recent biological motion simulation study also showed that low frequency components are critical for accurate speed estimation ^28^. Lappin, Tadin, Nyquist and Corn ^43^ showed that motion detection thresholds in low vision participants were similar to normally sighted participants for speeds > 2 degrees/second for foveal viewing. It is known that very coarse biological motion, using a few bright spots, can be sufficient for recognizing the human motion such as walking, running and dancing ^26^. Wood, Tyrrell and Carberry ^44^ showed that biological motion information helped drivers improve their detection of pedestrians in night driving tasks. As for collision avoidance, optical flow “would be sufficient for controlling braking”, according to Lee’s Tau theory^45,46^.

While, this study did not show an impact on HP performance due to the change in visual acuity, this finding does not necessarily suggest low visual acuity doesn’t have any negative impacts in the real world driving. At first look, the findings of this study might seem in contradiction of the findings of our previous study, which showed visually impaired people had lower HP scores than normally sighted ^24^. In addition to the difference between the participants groups involved in the two studies, the frame rate of visual stimuli was also a factor causing the difference. In the previous study, HP videos were presented in their original format, 25 Hz, while the frame rate in this study was 2.5 Hz. A lag caused by the slow frame rate in this study can lead to late detection of the potential hazards, but it may not have a major impact on the detection rate. Indeed, as we found, the detection rates of the two studies were not different, it was just the detection time of this study was longer than the previous study. We speculate that in the real world, normally sighted drivers may be able to spot hazards earlier than the visually impaired. It is just that the advantage of normally sighted drivers was compromised due to stimulus settings of this study. Nevertheless, it is unknown how dangerous a slightly later detection than normally sighted drivers would be for visually impaired drivers. It is possible that the driving safety of visually impaired drivers would not be compromised by this slightly delayed detection if they can compensate on other aspects, for instance, speed control.

Driving requires constant awareness of situation of the environment. While this study uses the HP paradigm, the same concept can be applied to situational awareness. Through motion perception, visually impaired drivers may be able to constantly “see” the road environment and traffic condition. However, logic would dictate that there must be visual acuity limit for safe driving, however further research is needed in order to identify this threshold., and the conclusion can be drawn that HP does not necessarily require high level of visual acuity. In addition, it is possible central vision loss due to some eye diseases, such as albino, may concur with affected motion perception function ^47^. The roles of motion perception and visual acuity, as well as the interaction between them, for driving safety require further investigation.

## Conclusion

The findings suggest that motion perception may play an important role in driving with impaired vision. The results of this study also showed that degrading visual acuity to 20/120 had very small effect on hazard perception in our experiment settings. Relatively, motion information appeared to have larger and significant effects. These findings might explain to some extent why there are many bioptic drivers who have been driving for years without involvement in crashes, even though they drive with low visual acuity most of the time.

## Acknowledgment

This work was supported in part by the NIH Grant R01 AG041974 and China Visiting Scholarship to XT.

